# Retinal Vessel Volume Density and Intraoperative Cortical Vascular Phenotypes in Moyamoya Disease

**DOI:** 10.64898/2026.11.19.767984

**Authors:** Liming Zhao, Shiliang Lou, Hao Liang, Alexandra Morozova, Yang Liu, Tao Gao, Yuxue Sun, Gaochao Guo, Ziqiang Liu, Zhenwei Li, Shangyu Jin, Ke Yang, Ke Li, Ivan Radovanovic, Hugo Andrade-Barazarte, Chaoyue Li

**Author notes:** Corresponding Author: Hugo Andrade-Barazarte MD PhD, 399 Bathurst Street, M5T2S8, Toronto, Canada.; Chaoyue Li MD PhD, Weiwu Road 7#, Jinshui District, Zhengzhou, 453000, China. Zhao Liming and Lou Shiliang contributed equally to this work.

## Abstract

**Background:** Angiographic staging describes large-artery stenosis and collateral evolution in moyamoya disease but does not directly depict the cortical surface vasculature, which can vary among patients with similar Suzuki stages. We examined whether a side-matched three-dimensional retinal vascular measure was associated with intraoperative cortical vascular phenotypes after accounting for Suzuki stage.

**Methods:** This prospective single-center cohort included 52 patients with moyamoya disease who underwent combined direct and indirect revascularization; 48 had analyzable retinal vessel volume density (RVVD) in the side-matched eye. RVVD was defined as the percentage of the corresponding retinal tissue volume occupied by three-dimensional optical coherence tomography angiography vascular signal within the central 6 × 6-mm region from the retinal nerve fiber layer to the inner plexiform layer. After dural opening and before vascular manipulation, overall cortical vascular richness and arterial, venous, and small-vessel richness were graded from 0 to 3. Associations were evaluated with Spearman correlation and proportional-odds ordinal logistic regression adjusted for age, sex, and ipsilateral Suzuki stage.

**Results:** Mean RVVD was 12.92% (SD, 3.30 percentage points). Higher RVVD was associated with greater overall cortical vascular richness (ρ=0.523; bias-corrected and accelerated bootstrap 95% CI, 0.279-0.714; P<0.001) and arterial richness (ρ=0.522; 95% CI, 0.261-0.714; P<0.001). Per 1-SD higher RVVD, the adjusted cumulative odds ratio for being in a higher richness category was 3.92 (95% CI, 1.82-8.44; P<0.001) for overall cortical vascular richness and 4.31 (95% CI, 1.92-9.70; P<0.001) for arterial richness.

**Conclusions:** In moyamoya disease, higher side-matched RVVD was associated with richer overall and arterial cortical vascular phenotypes at revascularization surgery, independent of Suzuki stage. Three-dimensional retinal vascular imaging may therefore provide a noninvasive, side-specific preoperative measure of cortical vascular remodeling.

**Nonstandard Abbreviations and Acronyms:** AVV, arterial vascular volume; AVVR, arterial-to-venous volume ratio; CFP, color fundus photography; DSA, digital subtraction angiography; MMD, moyamoya disease; OCT, optical coherence tomography; OCTA, optical coherence tomography angiography; RVVD, retinal vessel volume density; VVV, venous vascular volume.

## Introduction

Moyamoya disease (MMD) is a chronic, progressive occlusive cerebrovascular disorder that mainly affects the terminal internal carotid arteries and their proximal branches, with compensatory collateral formation and distal vascular remodeling [1–5]. In clinical practice, digital subtraction angiography (DSA) and Suzuki stage describe the evolution of large-artery stenosis, occlusion, and basal collateral networks, whereas perfusion imaging assesses the corresponding hemodynamic state. These examinations are central to diagnosis and treatment planning but do not directly depict the cortical surface vasculature seen after dural opening. During surgery, patients with similar Suzuki stages can show marked differences in pial arteries, veins, and small surface vessels. Previous studies have described varying degrees of cortical microvascularization and complex surface vascular remodeling in MMD [6–11].

The retina is one of the few neural microvascular beds that can be observed directly and noninvasively, and it shares some structural and physiological features with the cerebral microcirculation [12]. Retinal vascular signs have been associated with cerebrovascular disease and cerebral hemodynamic changes [13–16]. Optical coherence tomography (OCT) and optical coherence tomography angiography (OCTA) studies in MMD have identified retinal structural and microvascular abnormalities [17–20]. Hong et al. found that retinal photographs contained information relevant to MMD detection and angiographic staging [21]. However, whether these retinal vascular features correspond to the cortical surface vascular phenotype in the same patient remains unclear.

To address this question, we combined color fundus photography (CFP) with three-dimensional OCTA and OCT volume data to derive three-dimensional retinal vascular measures. The primary measure, retinal vessel volume density (RVVD), represents the proportion of the corresponding retinal tissue volume occupied by three-dimensional vascular signal in the macular region and retains volumetric information not represented by conventional two-dimensional vessel-density measures [22]. We examined whether RVVD in the side-matched eye was associated with cortical vascular richness directly observed during revascularization surgery and whether the association correlated to cerebral angiographic stage.

## Methods

### Study Design and Participants

This prospective, single-center observational study was conducted at the Moyamoya Disease Center of Henan Provincial People’s Hospital from November 2025 through April 2026. We consecutively screened patients older than 14 years whose MMD diagnosis was confirmed by digital substraction angiography (DSA) according to the 2021 revised diagnostic criteria [23]. Eligible patients underwent their first revascularization procedure, performed by the same surgeon. The operation combined superficial temporal artery-middle cerebral artery bypass with encephaloduro-myosynangiosis.

Exclusion criteria were severe cardiovascular or respiratory disease; Graves disease, antiphospholipid syndrome, systemic vasculitis, or Sjogren syndrome; ocular disease affecting media clarity or retinal image quality; severe cognitive impairment after stroke that precluded cooperation with imaging; or incomplete required clinical or imaging data. Of 126 patients screened, 74 were excluded under these criteria and 52 were enrolled. Four enrolled patients lacked analyzable RVVD in the side-matched eye because of source-data transfer or other technical data-availability issues; they remained in analyses that did not require RVVD. The study is reported in accordance with the STROBE statement [24].

The Ethics Committee of Henan Provincial People’s Hospital approved the study (approval No. 202605). The study was conducted in accordance with the Declaration of Helsinki. All participants or their legal representatives provided written informed consent.

Retinal Imaging and Three-Dimensional Vascular Quantification

Dedicated ophthalmic technicians performed all retinal examinations. A VG200D swept-source OCT/OCTA system was used to acquire fovea-centered 12 × 12-mm OCT and OCTA volumes. Color fundus photographs (CFP) with a 45-degree field of view were obtained using a ZEISS VISUCAM 224 fundus camera. .Image quality was assessed at the time of acquisition, with repeat imaging performed when necessary to obtain images of adequate quality.

Retinal measurements were matched to the surgical side: right-eye data were used for right-sided surgery and left-eye data for left-sided surgery. Retinal artery/vein segmentation and classification were automatically obtained from CFP using the image-processing pipeline used for this study. The resulting two-dimensional artery/vein labels were spatially registered to the en-face OCTA image and subsequently mapped into the three-dimensional OCT/OCTA volume. Vascular signals carrying arterial or venous labels were assigned to arterial vascular volume (AVV) or venous vascular volume (VVV), respectively. Capillaries and terminal small vessels that could not be classified reliably as arterial or venous were not assigned to AVV or VVV, but their signals remained part of the total vascular signal used to calculate RVVD (Figure 1).

**Figure 1.**
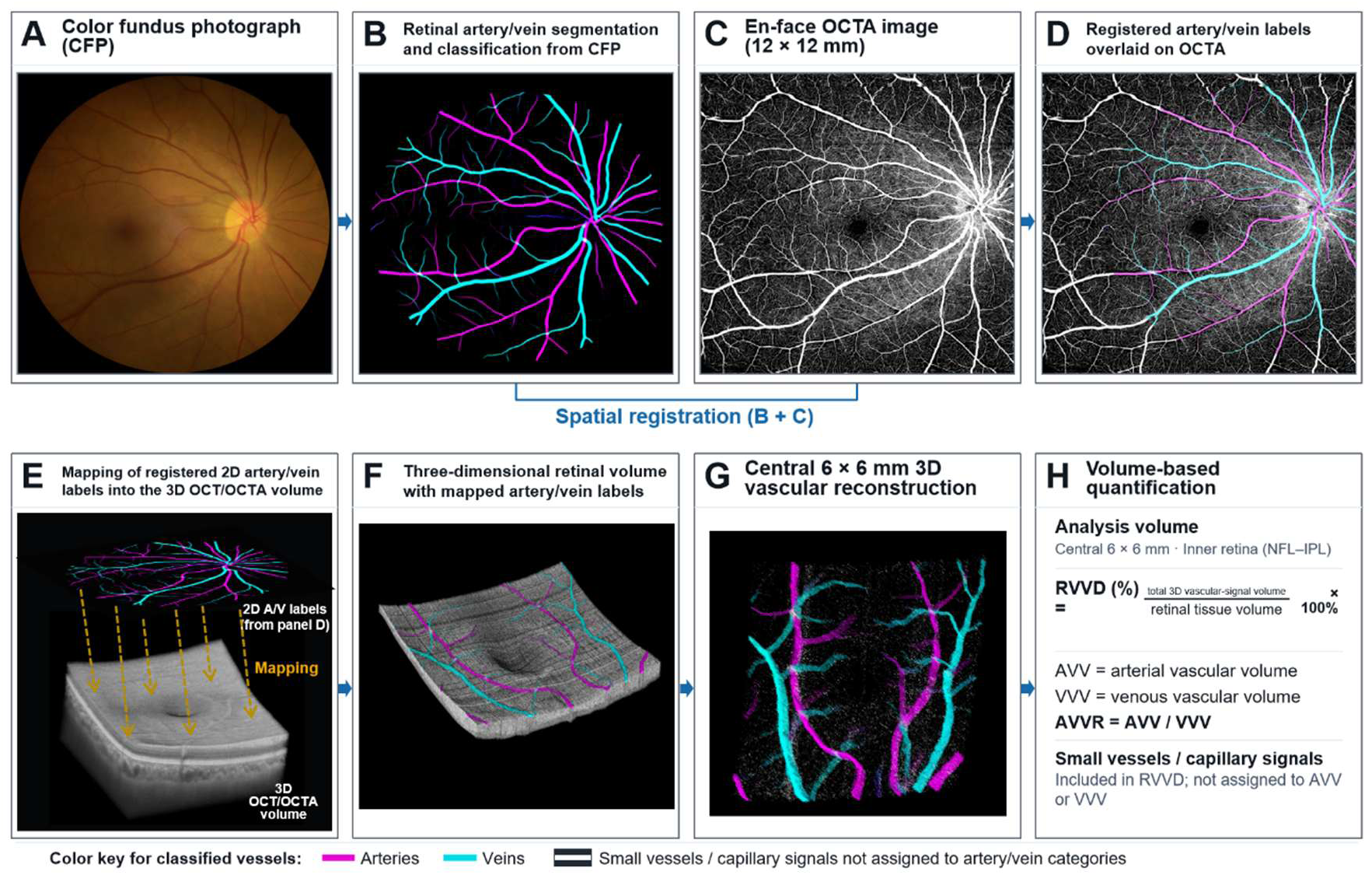
Multimodal retinal imaging and three-dimensional vascular quantification. (A) Color fundus photograph (CFP). (B) Retinal artery/vein segmentation and classification from the CFP image. (C) Fovea-centered 12 × 12-mm en-face optical coherence tomography angiography (OCTA) image. (D) Artery/vein labels after registration with the OCTA image. (E) Mapping of the registered two-dimensional labels into the three-dimensional OCT/OCTA volume. (F) Three-dimensional retinal volume with mapped artery/vein labels. (G) Three-dimensional vascular reconstruction of the central 6 × 6-mm region. (H) Volume-based quantification within the inner retina from the retinal nerve fiber layer to the inner plexiform layer (NFL–IPL). Retinal vessel volume density (RVVD) was calculated as total three-dimensional vascular-signal volume divided by retinal tissue volume ×100%. The arterial-to-venous volume ratio (AVVR) was calculated as arterial vascular volume (AVV) divided by venous vascular volume (VVV). Small-vessel and capillary signals not classified as arterial or venous were included in RVVD but not in AVV or VVV. Magenta indicates arteries, cyan indicates veins, and white indicates small-vessel or capillary signals. All retinal panels were obtained from the same patient and eye.

Using the center of the 12 × 12-mm scan, which corresponded to the foveal center, we directly cropped the central 6 × 6-mm region for analysis. Retinal layer segmentation was performed automatically by the device. The axial analysis range extended from the retinal nerve fiber layer to the inner plexiform layer. RVVD was defined as follows: RVVD (%) = total three-dimensional vascular-signal volume / corresponding retinal tissue volume ×100%.

The total vascular signal included detectable OCTA signals from large vessels, small vessels, and capillaries within this three-dimensional volume. RVVD is therefore an imaging-derived three-dimensional vascular volume fraction rather than a histologic measurement of true vascular lumen volume. The arterial-to-venous volume ratio (AVVR) was defined as AVV/VVV and was recorded as a dimensionless ratio without multiplication by 100. Foveal and parafoveal vessel volume densities, other retinal structural measures, and two-dimensional vascular measures were analyzed as exploratory variables. Two-dimensional arterial and venous densities were not interpreted as three-dimensional arterial or venous volume densities.

### Angiographic and Perfusion Assessment

Two neurosurgeons experienced in cerebrovascular angiography independently reviewed the DSA studies and resolved disagreements by consensus. Assessments included ipsilateral Suzuki stage, distal vascular-bed grade, posterior cerebral artery involvement, and leptomeningeal collateral grade. Regional cerebral blood flow and cerebral blood volume were obtained from routine magnetic resonance perfusion-weighted imaging. The clinical imaging system generated these measures from routine regions of interest; no additional study-specific postprocessing or manual adjustment was performed.

### Intraoperative Cortical Vascular Phenotype

Intraoperative photographs were obtained after dural opening and adequate exposure of the cortical surface but before bypass- or anastomosis-related vascular manipulation. The assessment covered the visible cortical surface within the craniotomy. All images were anonymized and renumbered before review.

Two neurosurgeons with at least 10 years of cerebrovascular surgical experience independently graded overall cortical vascular richness, arterial richness, venous richness, and small-vessel richness on a 0-to-3 scale. For overall, arterial, and venous richness, grade 0 indicated that the relevant vessels were almost absent or extremely sparse; grade 1, few vessels with large areas lacking visible vascular coverage; grade 2, a moderate number of identifiable vessels and branches without a dense network; and grade 3, abundant, densely branching vessels widely distributed across the operative field. For small-vessel richness, grade 0 indicated no or almost no small vessels; grade 1, a few scattered small vessels; grade 2, locally abundant small-vessel networks; and grade 3, a widespread, dense network of fine moyamoya-like surface vessels. Overall cortical vascular richness was a global visual assessment of the operative field and was not calculated as the sum of the arterial, venous, and small-vessel scores.

The raters were blinded to retinal measurements, angiographic and perfusion variables, postoperative status, and the other rater’s initial score. They jointly reviewed discrepant scores. If disagreement persisted, a third senior cerebrovascular neurosurgeon adjudicated the score. Consensus or adjudicated scores were used for analysis.

## Statistical Analysis

Continuous variables are reported as mean (SD) or median (interquartile range), according to their distributions, and categorical variables as number (percentage). The primary analyses examined RVVD in relation to overall cortical vascular richness and arterial richness. Venous richness and small-vessel richness were secondary outcomes.

Unadjusted associations between RVVD and the cortical vascular richness scores were evaluated with Spearman rank correlation. Bias-corrected and accelerated (BCa) bootstrap 95% CIs for Spearman ρ were estimated with 10,000 resamples [25]. RVVD across ordered score categories was summarized as median (interquartile range) and assessed with Cuzick’s Wilcoxon-type test for trend [26].

Because the intraoperative cortical vascular scores were ordinal, proportional-odds ordinal logistic regression was used to examine the association of RVVD with each phenotype. Results are reported as cumulative odds ratios (ORs) and 95% CIs for being in a higher richness category per 1-SD higher RVVD [27]. Multivariable models adjusted for age, sex, and ipsilateral Suzuki stage. Model convergence and multicollinearity were examined. The proportional-odds assumption was evaluated descriptively using binary models at different cumulative thresholds; sparse extreme score categories prevented a stable formal determination for some outcomes.

All retina-cortex comparisons other than the primary RVVD analyses for overall cortical vascular richness and arterial richness were exploratory. Benjamini-Hochberg correction was applied separately to the exploratory retina-cortex comparisons, angiographic/perfusion analyses [28]. Analyses used complete available cases without imputation. All tests were 2-sided, and P<0.05 was considered statistically significant. Analyses were performed with Python 3.12.

## Results

### Study Population

Of 126 patients screened, 74 were excluded according to the prespecified eligibility criteria and 52 were enrolled. Four enrolled patients lacked analyzable RVVD in the side-matched eye because of source-data transfer or other technical data-availability issues, leaving 48 patients in the primary RVVD analysis.

The median age of the primary cohort was 48 years (interquartile range, 41-54), and 22 patients (45.8%) were women. Thirty-seven (77.1%) had bilateral MMD, 42 (87.5%) had ischemic presentation, and 29 (60.4%) underwent left-sided surgery. Median ipsilateral Suzuki stage was 4 (interquartile range, 3-4); 4 patients had stage 2, 18 had stage 3, 15 had stage 4, 9 had stage 5, and 2 had stage 6 (Table 1).

**Table 1.**
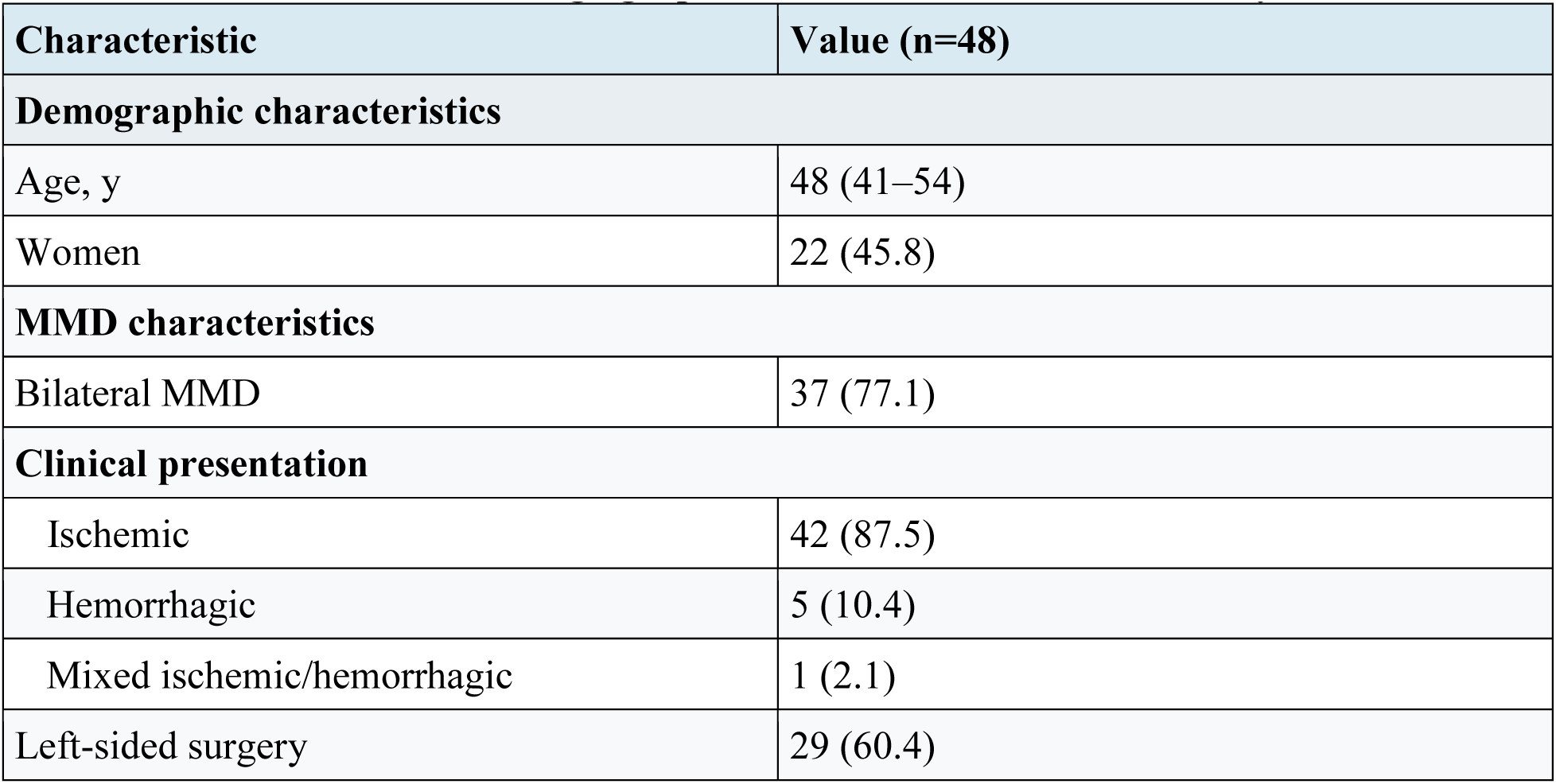
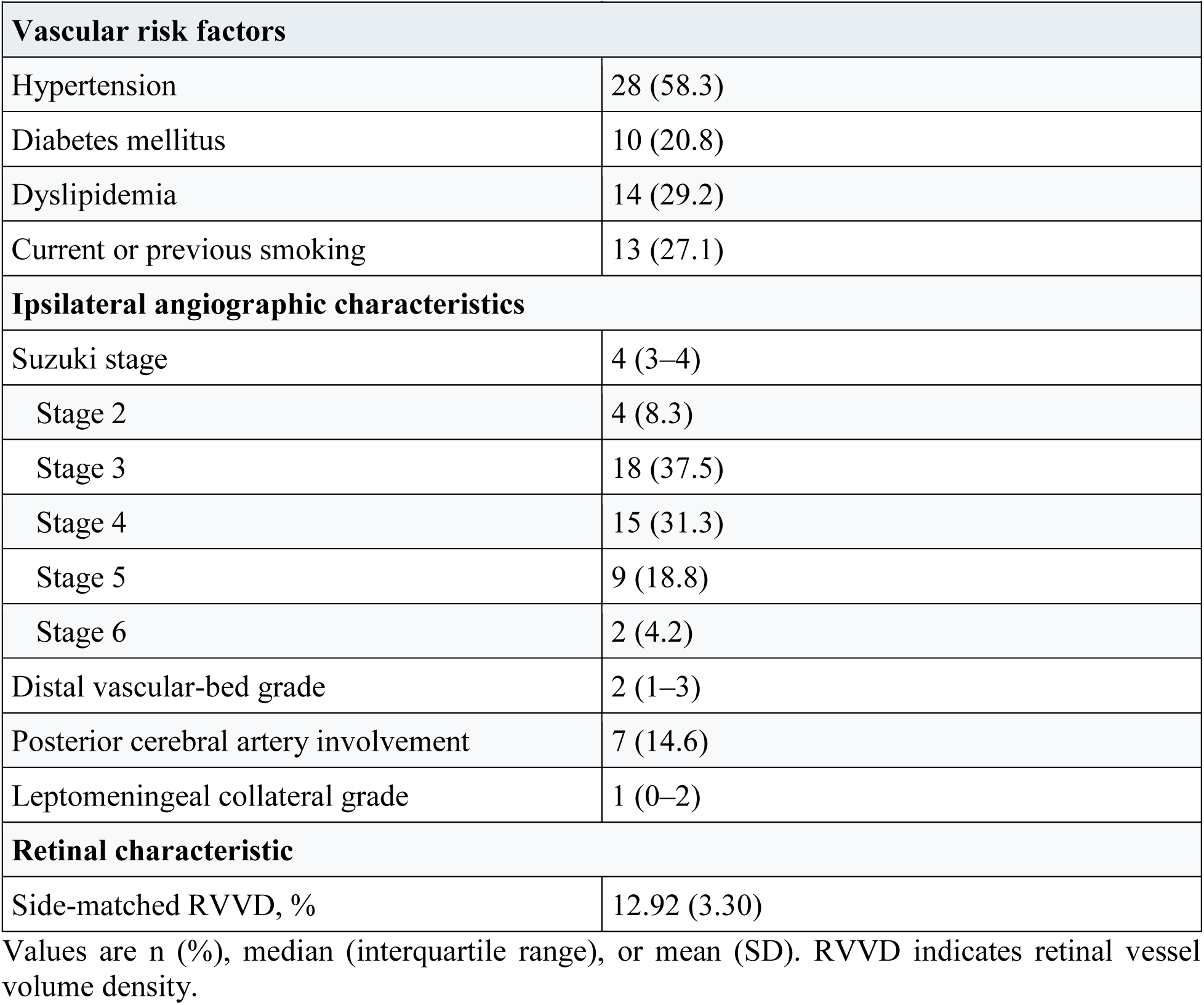
Baseline Clinical and Angiographic Characteristics of the Primary RVVD Cohort

Retinal Vascular Characteristics

Mean RVVD in the side-matched eye was 12.92% (SD, 3.30 percentage points), the median was 12.34% (interquartile range, 10.73%-14.38%), and the range was 3.72%-21.53%. RVVD, OCTA vessel density, and most regional three-dimensional or OCTA-derived measures were available in 48 patients. CFP-derived two-dimensional arterial and venous measures and AVVR were available in all 52 patients (Supplemental Table S1).

RVVD and the Intraoperative Cortical Vascular Phenotype

Higher RVVD was associated with greater overall cortical vascular richness and arterial richness. Representative side-matched retinal and intraoperative images across richness grades are shown in Figure 2. The Spearman correlation was 0.523 for overall cortical vascular richness (95% CI, 0.279-0.714; P<0.001) and 0.522 for arterial richness (95% CI, 0.261-0.714; P<0.001; Figure 3 and Supplemental Table S1).

**Figure 2.**
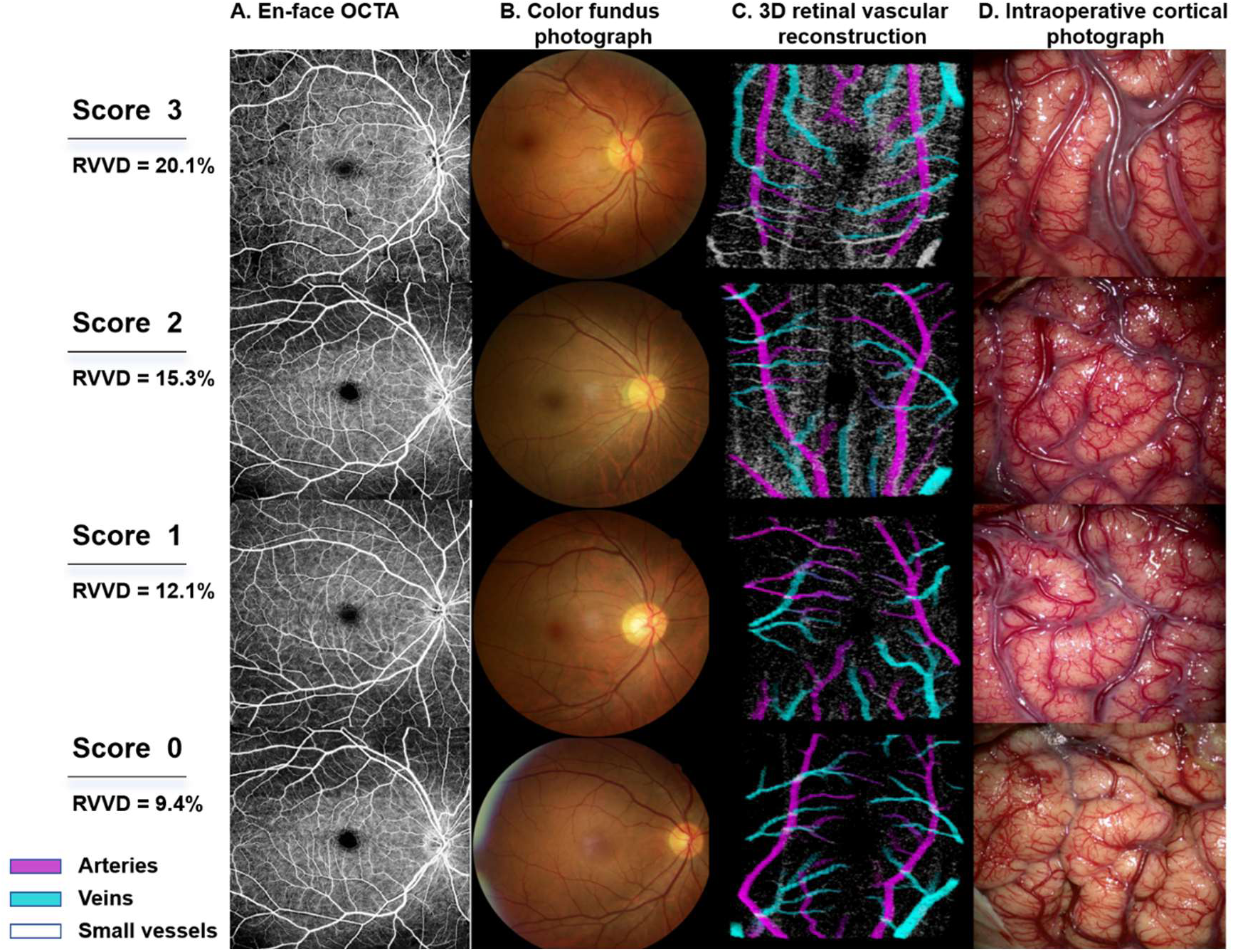
Representative retinal and intraoperative cortical images across different grades of cortical vascular richness. Representative cases are arranged from higher to lower overall cortical vascular richness scores (3 to 0), with corresponding RVVD values of 20.1%, 15.3%, 12.1%, and 9.4%. Each row shows the side-matched en-face OCTA image (A), color fundus photograph (B), three-dimensional retinal vascular reconstruction (C), and intraoperative cortical photograph obtained after dural opening (D) from the same patient. In the three-dimensional reconstructions, magenta indicates arteries, cyan indicates veins, and white indicates small-vessel or capillary signals. RVVD indicates retinal vessel volume density.

**Figure 3.**
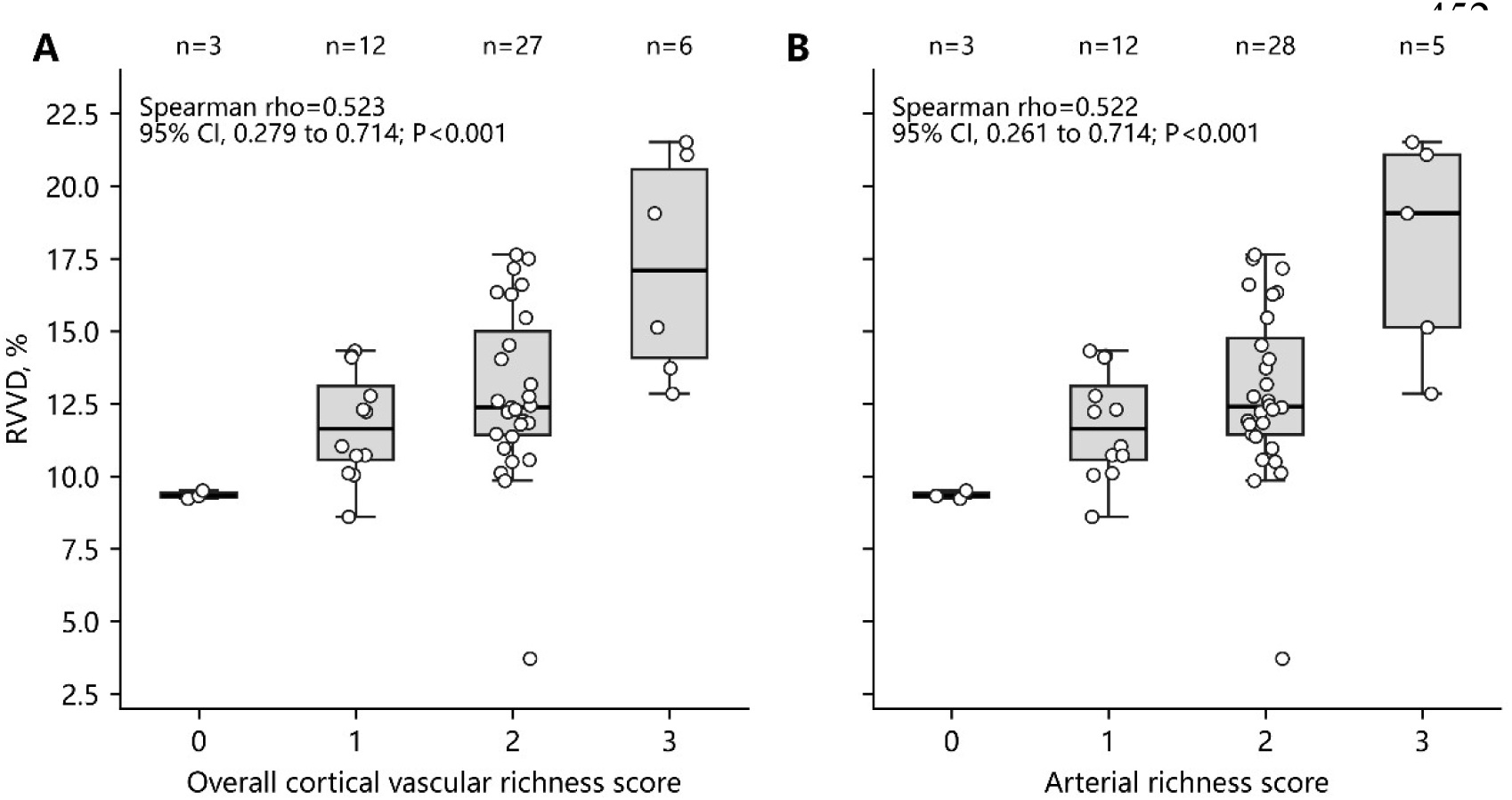
Retinal vessel volume density across grades of cortical vascular richness. Side-matched retinal vessel volume density (RVVD) is shown according to overall cortical vascular richness (A) and arterial richness (B), each scored from 0 to 3. Individual observations are shown together with box plots representing the median and interquartile range; whiskers extend to 1.5 times the interquartile range. Spearman correlation coefficients, 95% CIs obtained by bias-corrected and accelerated bootstrap resampling (10,000 replicates), and 2-sided P values are shown in each panel. Forty-eight patients with analyzable RVVD were included.

Across overall cortical vascular richness grades 0 through 3, median RVVD values were 9.33% (interquartile range, 9.28%-9.42%; n=3), 11.63% (10.57%-13.11%; n=12), 12.38% (11.42%-15.00%; n=27), and 17.10% (14.08%-20.58%; n=6), respectively (P for trend <0.001). Across arterial richness grades 0 through 3, median RVVD values were 9.33% (9.28%-9.42%; n=3), 11.63% (10.57%-13.11%; n=12), 12.41% (11.44%-14.76%; n=28), and 19.07% (15.13%-21.09%; n=5), respectively (P for trend <0.001).

Associations were weaker for the secondary phenotypes. The Spearman correlation was 0.317 for venous richness (95% CI, -0.009 to 0.583; P=0.028) and 0.299 for small-vessel richness (95% CI, -0.020 to 0.560; P=0.039); both bootstrap 95% CIs crossed zero (Supplemental Table S1).

## Multivariable Analysis

One SD of RVVD was 3.30 percentage points. After adjustment for age, sex, and ipsilateral Suzuki stage, the cumulative OR per 1-SD higher RVVD was 3.92 (95% CI, 1.82-8.44; P<0.001) for being in a higher overall cortical vascular richness category and 4.31 (95% CI, 1.92-9.70; P<0.001) for being in a higher arterial richness category (Table 2 and Figure 4). The corresponding adjusted ORs were 2.79 (95% CI, 1.31-5.94; P=0.008) for venous richness and 2.71 (95% CI, 1.32-5.60; P=0.007) for small-vessel richness.

**Figure 4.**
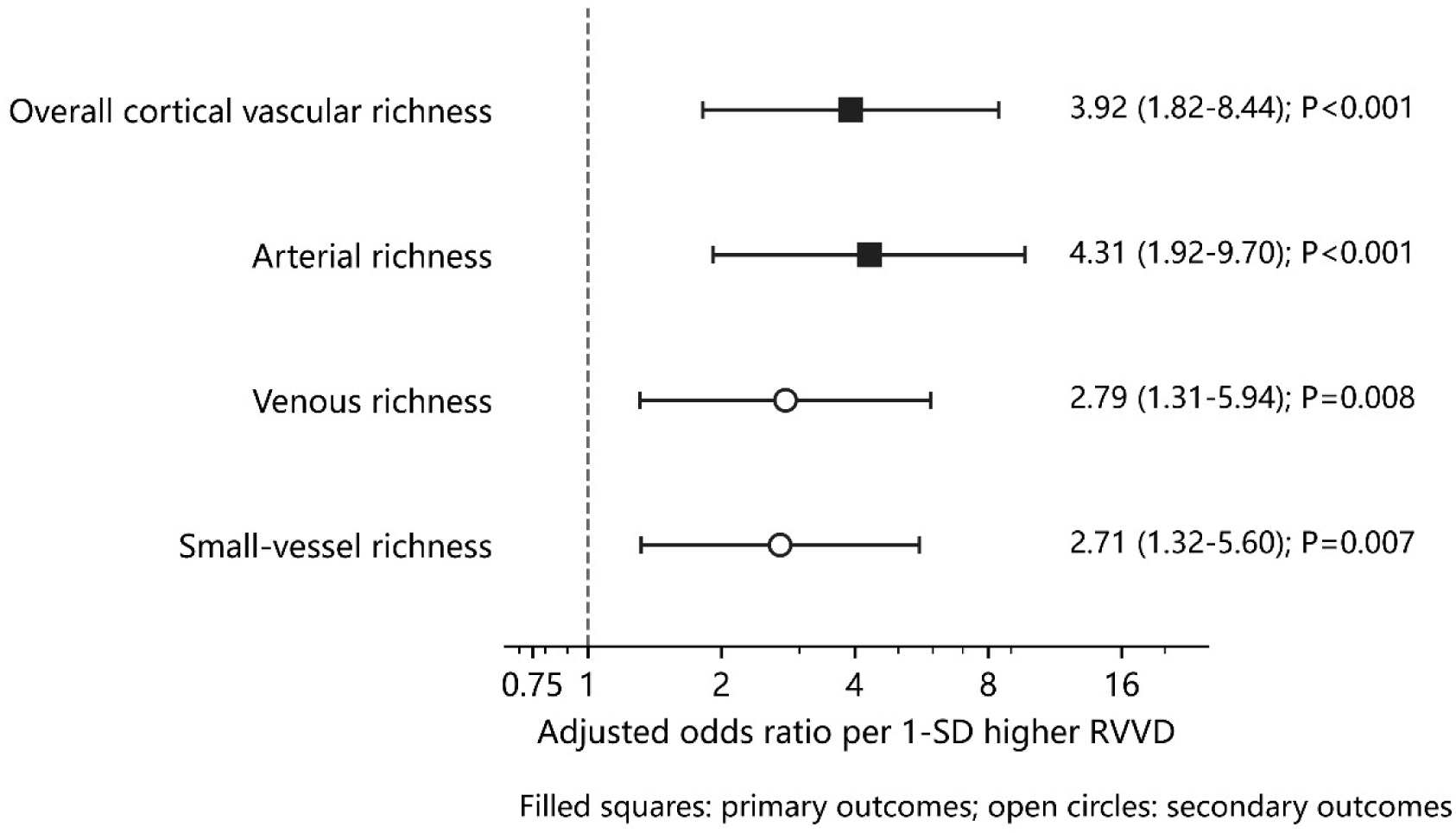
Adjusted associations between retinal vessel volume density and intraoperative cortical vascular phenotypes. Forest plots show cumulative odds ratios (ORs) and 95% CIs for being in a higher vascular richness category per 1-SD higher RVVD. Models were adjusted for age, sex, and ipsilateral Suzuki stage and included 48 patients. One SD of RVVD corresponded to 3.30 percentage points. Overall cortical vascular richness and arterial richness were the primary outcomes; venous and small-vessel richness were secondary outcomes. The vertical reference line indicates an OR of 1. RVVD indicates retinal vessel volume density.

**Table 2.** Multivariable Associations Between RVVD and Intraoperative Cortical Vascular Phenotypes

| **Outcome** | **n** | **Adjusted OR per 1-SD higher RVVD** | **95% CI** | **P value** | **Analysis level** |
|---|---|---|---|---|---|
| Overall cortical vascular richness | 48 | 3.92 | 1.82–8.44 | <0.001 | Primary |
| Arterial richness | 48 | 4.31 | 1.92–9.70 | <0.001 | Primary |
| Venous richness | 48 | 2.79 | 1.31–5.94 | 0.008 | Secondary |
| Small-vessel richness | 48 | 2.71 | 1.32–5.60 | 0.007 | Secondary |

All 4 proportional-odds models converged. Model diagnostics and the effect of sparse extreme score categories on assessment of the proportional-odds assumption are reported in the Supplemental Methods and Results.

Adjusted proportional-odds models included age, sex, ipsilateral Suzuki stage, and standardized RVVD. ORs are cumulative odds ratios for being in a higher outcome category. One SD of RVVD was 3.30 percentage points. Sparse extreme categories limited definitive formal assessment of the proportional-odds assumption for some outcomes; details are provided in the Supplemental Methods and Results. CI indicates confidence interval; OR, odds ratio; and RVVD, retinal vessel volume density.

Secondary and Exploratory Analyses

Foveal and parafoveal vessel volume densities were positively correlated with overall cortical vascular richness and arterial richness, with direction and magnitude similar to the RVVD findings. Associations of two-dimensional OCTA- and CFP-derived measures with the cortical phenotypes were less consistent, and AVVR showed no clear association. Complete results, including false-discovery-rate correction, are provided in Supplemental Table S1.

RVVD was weakly correlated with ipsilateral Suzuki stage (ρ=0.163; 95% CI, -0.129 to 0.423; P=0.267). No clear associations were found with distal vascular-bed grade (ρ=-0.239; 95% CI, - 0.478 to 0.056; P=0.102), posterior cerebral artery involvement (ρ=0.032; P=0.829), or leptomeningeal collateral grade (ρ=0.090; 95% CI, -0.195 to 0.365; P=0.542). Associations of RVVD showed no clear association with ipsilateral Suzuki stage or other angiographic measures. Associations with resting cerebral perfusion were also inconsistent and did not remain after correction for multiple comparisons (Supplemental Table S2). Thus, the observed retina–cortex association was not explained by a parallel association of RVVD with angiographic stage or resting perfusion.

## Discussion

The angiographic features of MMD include progressive intracranial arterial stenosis or occlusion and the development of abnormal collateral vessels, yet angiographic stage alone does not fully characterize the vascular phenotype. Cortical vascular richness varied among patients within the same Suzuki stage. RVVD was nevertheless associated with cortical vascular richness, and this association persisted after adjustment for ipsilateral Suzuki stage. This suggests that RVVD and Suzuki stage capture different aspects of the vascular phenotype. These associations persisted after adjustment for age, sex, and ipsilateral Suzuki stage. The retinal vascular phenotype obtained before surgery corresponded, to some extent, to the cortical vascular architecture directly observed during revascularization. The rationale for examining the retina in MMD patients, relies on the partial structural and physiological resemblance between retinal and cerebral microvascular beds [12]. Retinal vascular signs have been associated with cerebrovascular disease and cerebral hemodynamics [13–16], and OCT and OCTA studies have documented retinal structural and microvascular changes in MMD [17–20]. Hong et al. found that retinal photographs contained information relevant to MMD detection and angiographic stage [21]. This work indicates that MMD-related vascular information is present in retinal images, but it did not establish how the retinal phenotype relates to the cortical surface vasculature in the same patient.

The association of RVVD with overall cortical vascular richness and arterial richness may reflect the chronic vascular remodeling that characterizes MMD. Persistent restriction of arterial inflow may coexist with collateral recruitment, distal vascular adaptation, and changes in cortical microvascular structure [6–11]. Retinal and cortical vessels differ in their arterial supply and local regulatory environments and should not be regarded as anatomical equivalents. The present findings instead describe a phenotypic association between two neural vascular beds, with retinal vascular measurements potentially capturing structural features related to the cortical vascular state.

Intraoperative observations also showed that cortical vascular remodeling in MMD did not follow a single pattern. In some patients, larger recognizable cortical arteries and veins were attenuated or sparse, while numerous fine surface vessels produced a visibly red cortical surface densely covered by small vessels. In such cases, the richness of identifiable arteries and veins could be low despite abundant small vessels. Overall cortical vascular richness, arterial richness, venous richness, and small-vessel richness were related but represented different components of the cortical phenotype. The four intraoperative scores were designed to characterize different components of the cortical vascular phenotype rather than interchangeable measures of the same process. Associations of RVVD with venous and small-vessel richness were weaker in the unadjusted analyses and were therefore treated as secondary findings and interpreted cautiously. Volumetric quantification may have contributed to the observed retina-cortex relationship. Conventional CFP-derived or en-face OCTA vessel-density measures represent vascular signal after planar projection, which compresses depth information. RVVD quantifies the fraction of vascular signal within a defined retinal tissue volume and therefore retains some three-dimensional spatial information [22]. In this study, foveal and parafoveal vessel volume densities showed patterns similar to those for RVVD, whereas associations between planar measures and cortical vascular phenotypes were less consistent. The regional volume-based results were consistent with the principal RVVD result; the exploratory comparisons with planar measures do not establish comparative performance.

Each imaging modality addresses a different level of MMD pathology. Suzuki stage describes the angiographic evolution of large-artery stenosis or occlusion and basal collateral networks, whereas other DSA variables characterize specific collateral routes or the distal vascular bed. Transdural collateral flow has been associated with hemodynamic differences in adults with MMD [29]. Perfusion imaging measures tissue hemodynamics at the time of acquisition; prior computed tomography perfusion studies have reported stage-related hemodynamic differences in ischemic MMD [30]. Intraoperative photographs showed cortical surface vascular morphology, whereas RVVD quantifies three-dimensional vascular signal in the retinal neurovascular bed. In the present study, the weak or inconsistent associations of RVVD with Suzuki stage and resting perfusion did not conflict with its association with the cortical surface phenotype. Persistence of the association after adjustment for ipsilateral Suzuki stage is consistent with RVVD capturing information not fully represented by angiographic stage.

This study has several limitations. It was conducted at a single center, included a modest sample of patients with MMD, and lacked independent external validation. The intraoperative cortical phenotypes were graded semiquantitatively and may have been influenced by the extent of surgical exposure and visual judgment, although experienced cerebrovascular neurosurgeons scored the images independently while blinded to other study variables. The cross-sectional, associational design cannot determine the biological direction of the retina-cortex relationship. RVVD was derived from OCTA vascular signal and multimodal three-dimensional reconstruction; its repeatability, registration performance, and transferability across devices require further assessment. Four enrolled patients did not have analyzable side-matched RVVD, and the number of postoperative events was too small to support firm conclusions about early clinical outcomes. Multicenter studies should incorporate repeatability assessment, quantitative analysis of cortical surface vasculature, and longitudinal retinal imaging.

## Conclusions

In patients with MMD, higher RVVD in the side-matched eye was associated with greater overall cortical vascular richness and arterial richness observed during surgery. The associations persisted after adjustment for ipsilateral Suzuki stage. Three-dimensional retinal vascular imaging may reflect aspects of cortical vascular remodeling that are not directly represented by conventional angiographic staging.

## Data Availability

Deidentified data that support the findings of this study may be available from the corresponding author on reasonable request, subject to institutional approval and applicable ethical and privacy requirements. Individual-level source data are not deposited in a public repository.

## Ethics Statement

The study was approved by the Ethics Committee of Henan Provincial People’s Hospital (approval No. 202605) and was conducted in accordance with the Declaration of Helsinki. Written informed consent was obtained from all participants or their legal representatives.

## Author Contributions

Liming Zhao contributed to conceptualization, methodology, and writing of the original draft. Shiliang Lou contributed to methodology, visualization, and data curation. Hao Liang contributed to investigation and data curation. Alexandra Morozova contributed to writing - review and editing. Yang Liu, Tao Gao, and Yuxue Sun contributed to data curation. Gaochao Guo and Ziqiang Liu contributed to formal analysis. Zhenwei Li, Shangyu Jin,Ke Yang and Ke Li contributed to investigation and data curation. Hugo Andrade-Barazarte contributed to conceptualization, supervision, and writing - review and editing. Chaoyue Li contributed to resources, funding acquisition, and project administration. Ivan Radovanovic contributed to supervision and writing - review and editing. All authors reviewed and approved the final manuscript.

## Data Access and Responsibility

Liming Zhao had full access to all source data and take responsibility for the integrity of the data, the analysis, and the accuracy of the statistical results.

